# A new approach to automated CBMN scoring following high doses

**DOI:** 10.1101/620971

**Authors:** Mikhail Repin, David J. Brenner, Guy Garty

## Abstract

In recent years we have automated the CBMN assay using microvolumes of blood, processed in multiwell plates. We have seen that at doses above 6 Gy the detected yield of micronuclei actually declines with dose, likely because of mitotic delay, preventing cells from forming micronuclei and also, when using one color imaging, resulting in many false binucleated cells, consisting of two randomly-adjacent nuclei. By using the inverse mitotic index (the ratio of mononuclear to binuclear cells) to adjust the micronucleus yield we were able to obtain a monotonic increasing dose response curve at doses of up to at least 10 Gy from the same samples which generated dose-response curve with a peak near 6 Gy, when scored using the traditional micronucleus yield.

## Introduction

Following a large scale radiological event there will be an immediate need to locate individuals who were exposed to high doses and would benefit from medical treatment. Over the past years the Center for High Throughput Minimally Invasive Radiation Biodosimetry at Columbia University has focused on development of robotic platforms for high throughput processing of cytogenetic bioassays like the Cytokinesis Blocked Micronucleus assay (CBMN)^(1)^ and the Dicentric Chromosome Assay (DCA). The Robotic Automated Biodosimetry Technology (RABiT II)^(2)^ we have developed makes use of commercially available High Throughput Screening platforms to perform these assays in 96-well microplates, using 20-30 μl of whole blood per sample.

In the traditional CBMN assay^(3, 4)^ lymphocytes are cultured to division and cytokinesis is blocked, forming binucleated cells. Chromosomes or chromosomal fragments which were damaged by the radiation are ejected from the daughter nuclei and manifest as micronuclei within the same cytoplasm, with the average number of micronuclei per binucleated cell (Mni/Bi)increasing with dose.

A major issue we^(5)^, and others^(6)^ have encountered with the CBMN assay is a decreasing yield of micronuclei at high doses, resulting in essentially the same assay readout for low and high doses and preventing the assay from being reliably used without further knowledge.

Using the conventional micronucleus assay, a few ml of blood is cultured and several microscope slides prepared and manually scored per sample. In this case micronucleus yields are monotonically increasing up to about 10 Gy^(6)^ – a different endpoint must be used for dose reconstruction if a high dose is suspected.

This problem becomes more severe when performing automated scoring. When scoring manually, the cells are typically stained with Giemsa and the cellular boundaries are obvious. Automated scoring more commonly utilizes DAPI which only stains nuclei. While it is possible to add a second stain to identify the boundaries of each cell, this is rarely done, as it significantly reduces throughput. More commonly, nuclei are associated into cells by proximity. In our prior work^(5)^ we developed an advanced algorithm for efficient detection of micronuclei, using high resolution (40x objective lens) imaging of sparsely-plated lymphocytes on microscope slides prepared from 1 ml blood samples. In that work, the turnover was observed at doses of 6-8 Gy, depending on the donor. Simultaneously, we had seen that the ratio of mononucleated to binucleated cells (*Mo/Bi*; related to the mitotic index) is monotonically increasing at high doses but fairly constant at low doses. Our solution was to implement a two-step scoring procedure where if *Mo/Bi* is large it is used for dose reconstruction, otherwise the conventional *Mni/Bi* is used.

More recently, we have developed a fully automated micronucleus assay, starting from 20-30 μl of whole blood, cultured and imaged at lower resolution (20x) in 96-well plates^(1, 7)^ with a fully automated image analysis routine. As can be seen below, the ratio *Mni/Bi,* traditionally used as a biomarker, reproduces the previously observed turnover at a dose of 6 Gy using these analysis conditions, and thus cannot be used for dose prediction of higher doses.

We present here a modified phenomenological micronucleus parameter that utilizes a combination of the traditional micronucleus yield and the mitotic index so that it is monotonically increasing with dose up to at least 10 Gy, using our microculture assay, and is thus useful as a biomarker for high, survivable, doses.

## Materials and Methods

Following informed consent (Columbia Institutional Review Board protocol # AAAF2671), blood from 8 volunteers was obtained by venipuncture. The blood was split into 20 μl aliquots and two aliquots per donor were irradiated to each of 0, 2, 4, 6, 8, 10 Gy, using a Gammacell 40 ^137^Cs irradiator (Atomic Energy of Canada Ltd, Chalk River, ON, Canada) at a dose rate of 0.7 Gy/min.

### Automated CBMN Assay

The Automated CBMN assay was performed as described in^(1)^ using a cell::explorer system (PerkinElmer, Waltham, MA). Briefly, PB-MAX karyotyping medium (Thermo Fisher Scientific Inc., Waltham, MA) was added to the samples in 96-well plate for a total volume of 250 μl/well. All samples were placed in the incubator at 37°C with 5% CO2. After 44 h of incubation, cytochalasin-B (Thermo Fisher) was added to the final concentration of 6 μg/ml and samples were cultured for another 26 h. After completion of culturing, cells were fixed and DAPI-stained as described before^(1)^.

### Automated Imaging

Prepared imaging microplates were scanned on an IN Cell Analyzer 2000 system^(8)^ (General Electric, Boston, MA). This automated imager is equipped with a large sensor CCD camera (2048 x 2048 pixel resolution) that is capable of whole-well imaging in 96- or 384-well microplates at high sensitivity. For this work, eighty one 20x fields, covering an area of 60 mm^2^ were captured per well, using an exposure of 300 msec/image.

### Automated Image Analysis

Image analysis was performed by using our custom software FluorQuant (v6.1)^(9)^, written in Visual C++ using the OpenCV computer vision libraries (Version 3.1, www.opencv.org). The software accepts a set of 16 bit grayscale images, directly from the RABiT II imaging system. The images are reduced to 8 bit by locating the brightest pixel value, *V,* in the image and dividing all other pixels by *f=V/255.* They are then binarized using an adaptive threshold algorithm, which assigns each pixel a value of 1 if its value is larger than pixels in a 25×25 pixel neighborhood and zero otherwise. Nuclei are then located as Binary Large Objects (BLOBs), using the algorithm of Suzuki and Abe^(10)^. Round BLOBS within a size range of 500-1000 pixels (corresponding to 70-140 μm^2^) are identified as nuclei and grouped into cells based on proximity – if two similarly sized nuclei are within 3 radii of each other, they belong to the same cell. Cells with two nuclei having dissimilar size (20% difference) or brightness (5% difference) are scored as two mononucleated cells, rather than as a binucleated cell. Micronuclei are identified as BLOBS, within the inferred cell, having an area 1/5 to 1/256 of the area of the main nuclei. The yield of Mononucleated, Binucleated, Trinucleatedr and Quadrinucleated cells and the total number of micronuclei in each category are reported in a Microsoft Excel file. Yields from the two aliquots were summed and plotted below.

Figure 1 shows the yield of micronuclei per binucleated cell in blood samples from 8 donors irradiated at doses of up to 10 Gy after fully automated image analysis. It is evident that the yield of micronuclei is a good biomarker for doses up to about 6 Gy. At higher doses the detected yield of micronuclei declines and a dose reconstruction would yield an erroneously low dose.

**Figure 1:**
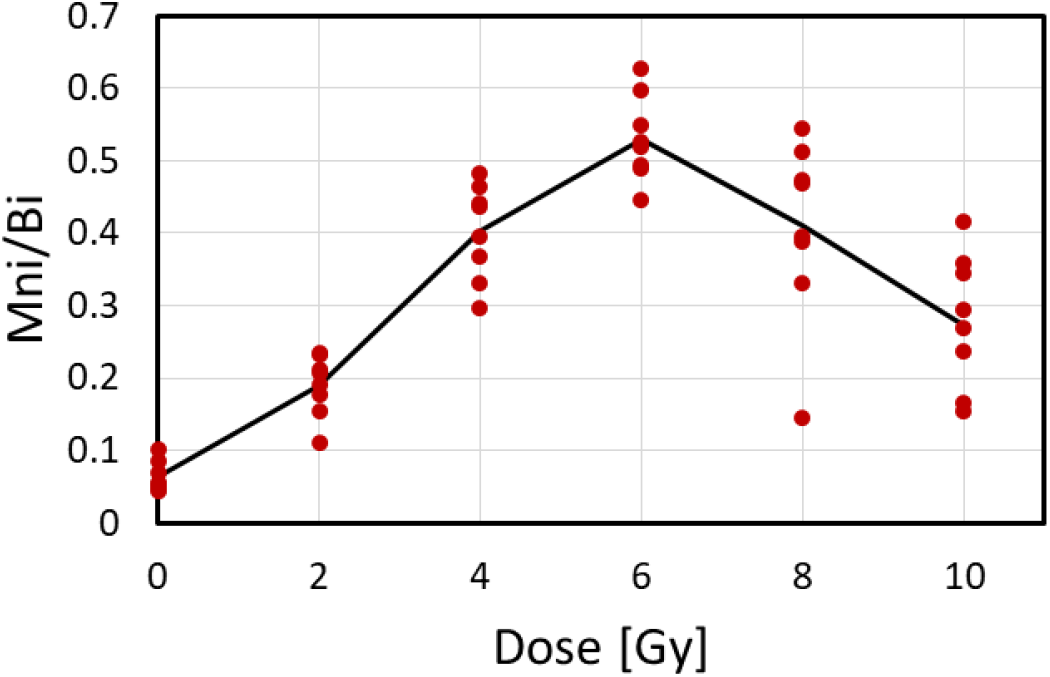
Yield of micronuclei per binucleated cell in irradiated samples from 8 healthy donors. The dots correspond to the yield scored for each donor. The line is the average of all samples, processed concurrently, in the same 96-well plate.

Figure 2 shows the ratio of the mononuclear cells to binucleated cells. This marker provides a good dose response at high doses but is generally considered to perform poorly at low doses^(5)^, where it is almost flat and suffers large variations between donors.

**Figure 2:**
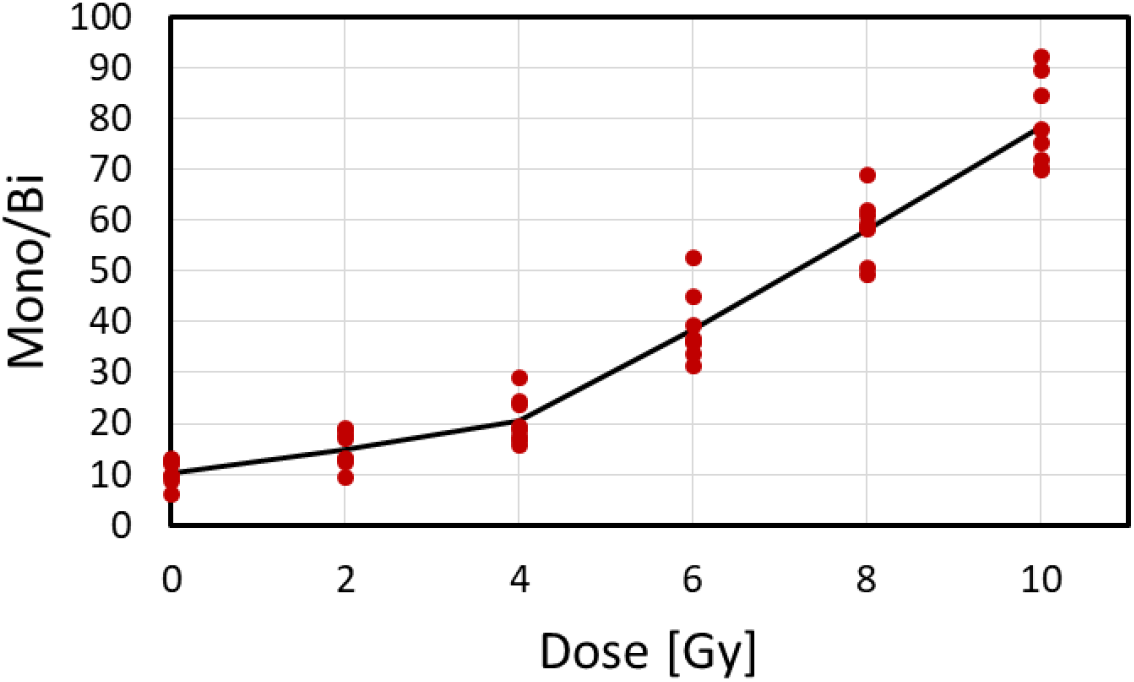
Ratio of mononucleated to binucleated cells in irradiated samples from 8 healthy donors. The dots correspond to the yield scored for each donor. The line is the average of all samples, processed concurrently, in the same 96-well plate.

## Discussion

Many labs use semi-automated imaging to perform the micronucleus assay^(11)^. Briefly, an automated microscope acquires images of cells from a slide and classifies them into binucleated cells and binucleated cells with 1 or more micronuclei, providing a gallery of images of candidate cells. A trained scorer then reviews the gallery and rejects or reclassifies the incorrectly classified cells. This is obviously a slow process limiting the number of samples scored. Once the misclassified cells are removed the measured micronucleus yield increases^(12)^ and the high dose turnover in the curve is reduced or eliminated.

It is extremely difficult to train software to perform this screening automatically and usually results in either insufficient rejection of misclassified cells or over-rejection and poor statistics. This is particularly severe when performing the assay in microculture, where there are not many binucleated cells to begin with. Another problem at high doses is that the yield of Binucleated cells decreases, mainly due to mitotic delay. Thus the sample contains many mononucleate cells which, due to the high density plating, may be misidentified as binucleated cells.

In prior work^(5)^ we have seen that the ratio of mononuclear (undivided) to binucleated cells is a good biomarker at high doses (above 3-4 Gy) at lower doses the response tends to be flat and there is large variation between individuals, although the use of better culture media seems to reduce this variation. The goal of combining these two markers into a single unified parameter is to get response similar to the yield of micronuclei at low doses and yet monotonically increasing.

A simple way to do that is to provide a weighted sum of the two, 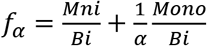, with α set such that the first term dominates at low doses and the second term dominates at high doses. Ideally we would also want *f* to be linear with dose, which simplifies dose reconstruction. By fitting *f* to a linear curve, using the built in routines in Microsoft Excel, we found that the best *R*^2^ was obtained for an α value of 45±5.

**Figure 3:**
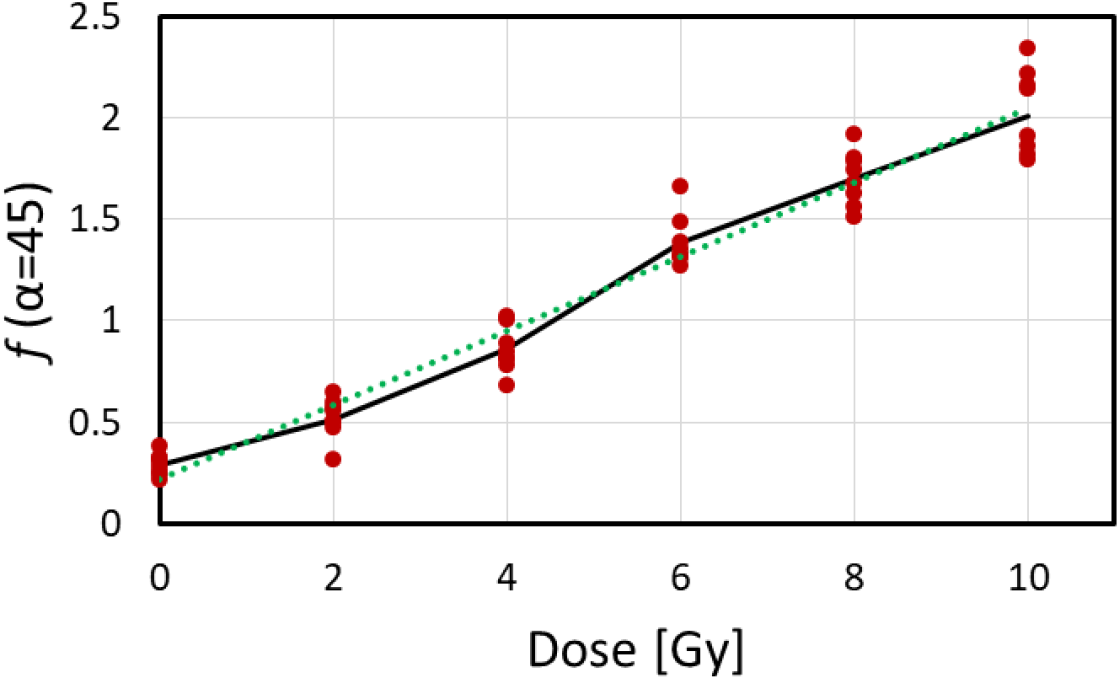
Dose dependence of f_45_(see text) in irradiated samples from 8 healthy donors. The dots correspond to the yield scored for each donor. The line is the average of all samples, processed concurrently, in the same 96-well plate.

## Summary

We present a novel approach, based on the use of phenomenological parameter – the classical yield of micronuclei corrected by mitotic index component – as a dose indicator. The new parameter is characterized by a linear dose response curve in a wide dose range from 0 up to 10 Gy and can be used for automated high dose prediction based on the automated micronucleus assay in microculture, where the more traditional micronucleus yield fails. By using mitotic index to correct the yield of micronuclei, a near linear dose response curve can be obtained. Thus it is possible to identify individuals who were exposed to higher doses and provide the appropriate care.

Considering that in most radiological scenarios^(13)^, the majority of individuals seeking treatment will either not be exposed or will have received low doses (“Worried Well”). It is thus crucial to be able to separate individuals who received a high dose from the rest of the population without requiring a trained specialist to review a gallery of micronucleus images for each individual, as is currently recommended. By using our approach, the dose can either be predicted automatically or a gallery of cell images can be generated for review by a specialist only for those individuals suspected of having received a high dose.

## Acknowledgments

This work was supported by grant number U19 AI067773, the Center for High-Throughput Minimally-Invasive Radiation Biodosimetry, from the National Institute of Allergy and Infectious Diseases, National Institutes of Health. The content is solely the responsibility of the authors and does not necessarily represent the official views of National Institute of Allergy and Infectious Diseases of the National Institutes of Health.

## References

1. Repin, M., Pampou, S., Karan, C., Brenner, D. J. and Garty, G. RABiT-II: Implementation of a High-Throughput Micronucleus Biodosimetry Assay on Commercial Biotech Robotic Systems. Radiat Res. 187(4), 502–508 2017.

2. Repin, M., Turner, H. C., Garty, G. and Brenner, D. J. Next generation platforms for high-throughput biodosimetry. Radiat Prot Dosimetry. 159(1-4), 105–110 2014.

3. Fenech, M. Cytokinesis-block micronucleus cytome assay. Nat Protoc. 2(5), 1084–1104 2007.

4. International Atomic Energy Agency. Cytogenetic dosimetry: applications in preparedness for and response to radiation emergencies. IAEA Emergency Preparedness and Response Series. IAEA 2011. IAEA 2011.

5. Lyulko, O. V., Garty, G., Randers-Pehrson, G., Turner, H. C., Szolc, B. and Brenner, D. J. Fast image analysis for the micronucleus assay in a fully automated high-throughput biodosimetry system. Radiat Res. 181(2), 146–61 2014.

6. Muller, W. U. and Rode, A. The micronucleus assay in human lymphocytes after high radiation doses (5-15 Gy). Mutat Res-Fund Mol M. 502(1-2), 47–51 2002.

7. Repin, M., Pampou, S., Garty, G. and Brenner, D. J. RABiT-II: A Fully-Automated Micronucleus Assay System with Shortened Time to Result. Radiat Res. 191(3), 232–236 2019.

8. Radu, C., Adrar, H. S., Alamir, A., Hatherley, I., Trinh, T. and Djaballah, H. Designs and concept reliance of a fully automated high-content screening platform. J Lab Autom. 17(5), 359–69 2012.

9. Garty, G., Bigelow, A. W., Repin, M., Turner, H. C., Bian, D., Balajee, A. S., Lyulko, O. V., Taveras, M., Yao, Y. L. and Brenner, D. J. An Automated Imaging System for Radiation Biodosimetry. Microsc Res Tech. 78(7), 587–598 2015.

10. Suzuki, S. and Abe, K. Topological structural analysis of digitized binary images by border following. Computer Vision, Graphics, and Image Processing. 29(3), 396 1985.

11. Depuydt, J., Baeyens, A., Barnard, S., Beinke, C., Benedek, A., Beukes, P., Buraczewska, I., Darroudi, F., De Sanctis, S., Dominguez, I. et al. RENEB intercomparison exercises analyzing micronuclei (Cytokinesis-block Micronucleus Assay). Int J Radiat Biol. 93(1), 36–47 2017.

12. Romm, H., Barnard, S., Boulay-Greene, H., De Amicis, A., De Sanctis, S., Franco, M., Herodin, F., Jones, A., Kulka, U., Lista, F. et al. Laboratory intercomparison of the cytokinesis-block micronucleus assay. Radiat Res. 180(2), 120–8 2013.

13. Homeland Security Council. National planning scenarios (final version 21.3) 2006.

